# Development of py-mea-span for comprehensive analysis of multi-electrode array data and its application to LGI2-deficient neurons

**DOI:** 10.64898/2026.10.14.759118

**Authors:** Zhuang Z. Han, Janet E. Deane

## Abstract

**Summary:** Multi-electrode arrays (MEAs) generate mid- to high-throughput electrophysiological data widely used to characterise neural network activity in vitro. Downstream analysis of recordings from Axion Biosystems instruments, however, has long depended on the manufacturer’s closed-source pipeline (AxIS Navigator, NeuralMetric Tool, AxIS Metric Plotting Tool), limiting customisation and integration into flexible Python workflows. We present py-mea-span, an open-source Python package providing end-to-end analysis of Axion Biosystems spk binary files. Our package enables spike extraction, inter-spike interval burst detection, network burst detection and spike synchrony. We also provide tools for preparation of publication-ready visualisations including raster plots with spike-detection-rate histograms, plate-layout heatmaps, time-course trajectories, single timepoint violin plots and statistical comparisons. Outputs reproduce the manufacturer’s NeuralMetric Tool reference with near perfect accuracy in all benchmarked metrics. A mea-span command-line interface enables batch processing without writing Python code. As a proof of concept, we applied py-mea-span to human iPSC-derived cortical neurons with CRISPRi-mediated LGI2 knockdown. This revealed a reproducible network hyperexcitability phenotype in which burst and network burst frequencies were elevated while spike synchrony remained unchanged.

**Availability and implementation:** The package is distributed on PyPI (pip install py-mea-span), and version-controlled at https://github.com/Molecularbiologyworld/py-mea-span. User feedback can be provided via the GitHub issues tab.

## INTRODUCTION

Multi-electrode arrays (MEAs) are now widely used to record spontaneous and evoked extracellular activity from cultured neuronal networks, enabling functional characterisation through metrics such as mean firing rate, burst frequency, network burst frequency, and inter-electrode synchrony ^1^. Recordings are non-invasive, so the same cultures can be followed from network formation to maturity. Multi-well plate formats make this tractable at screening scale. This approach now spans a broad range of applications. Human iPSC-derived neurons are used to model epilepsy-associated channelopathies ^2^, amyotrophic lateral sclerosis ^3^, and autism-spectrum disorders ^4^. MEA also underpins drug seizurogenicity ^5^ and neurotoxicity screening ^6^. Additionally, they support longitudinal studies of network maturation in monolayers ^7^ and in 3D organoids ^8^.

Among commercial MEA platforms, instruments from Axion Biosystems (the Maestro series) are widely deployed and store recordings in a proprietary spk binary format. Downstream analysis of these recordings has depended on the manufacturer’s closed-source pipeline, such as the NeuralMetric Tool, which limits flexibility and integration into custom analytical workflows and publication-ready visualizations.

A growing ecosystem of open-source tools address parts of the MEA analysis pipeline (**Table 1**), including MEAanalysis ^9^, meaRtools ^10^, and MEA-ToolBox ^11^. However, none of them directly reads the Axion spk binary format: each requires that the user first run NeuralMetric Tool and export tabular summaries, re-introducing a laborious manual step into otherwise automated workflows.

**Table 1.**
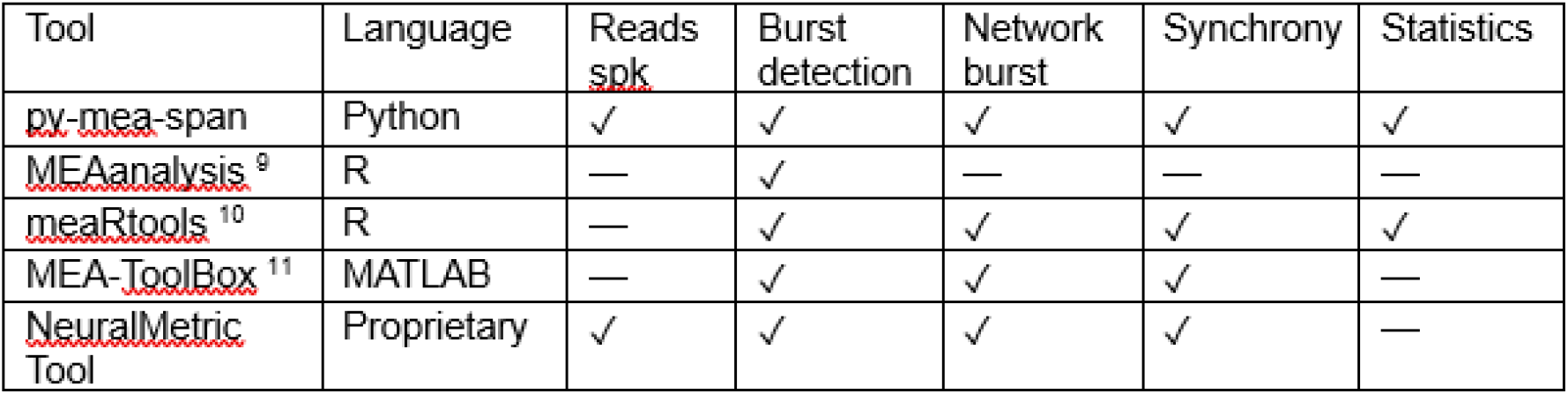
Comparison of py-mea-span with a selected subset of existing tools for MEA data analysis.

Here we present py-mea-span (Python Multi-Electrode Array Signal Processing and ANalysis), an open-source Python package that closes this gap. It directly parses the spk files, and computes the full set of single-electrode and network-level metrics in very close agreement with the manufacturer’s NeuralMetric Tool reference. The package extracts and analyzes raw data as well as providing a range of options for data visualization and statistical comparisons. The package can be used in two ways: by importing it into a Python script for custom analysis, or by typing mea-span in the command line interface (CLI) in a terminal. The latter option requires no Python code, only an installed Python environment. Twenty-seven metrics (**Table 2**) are covered in this package.

**Table 2.**
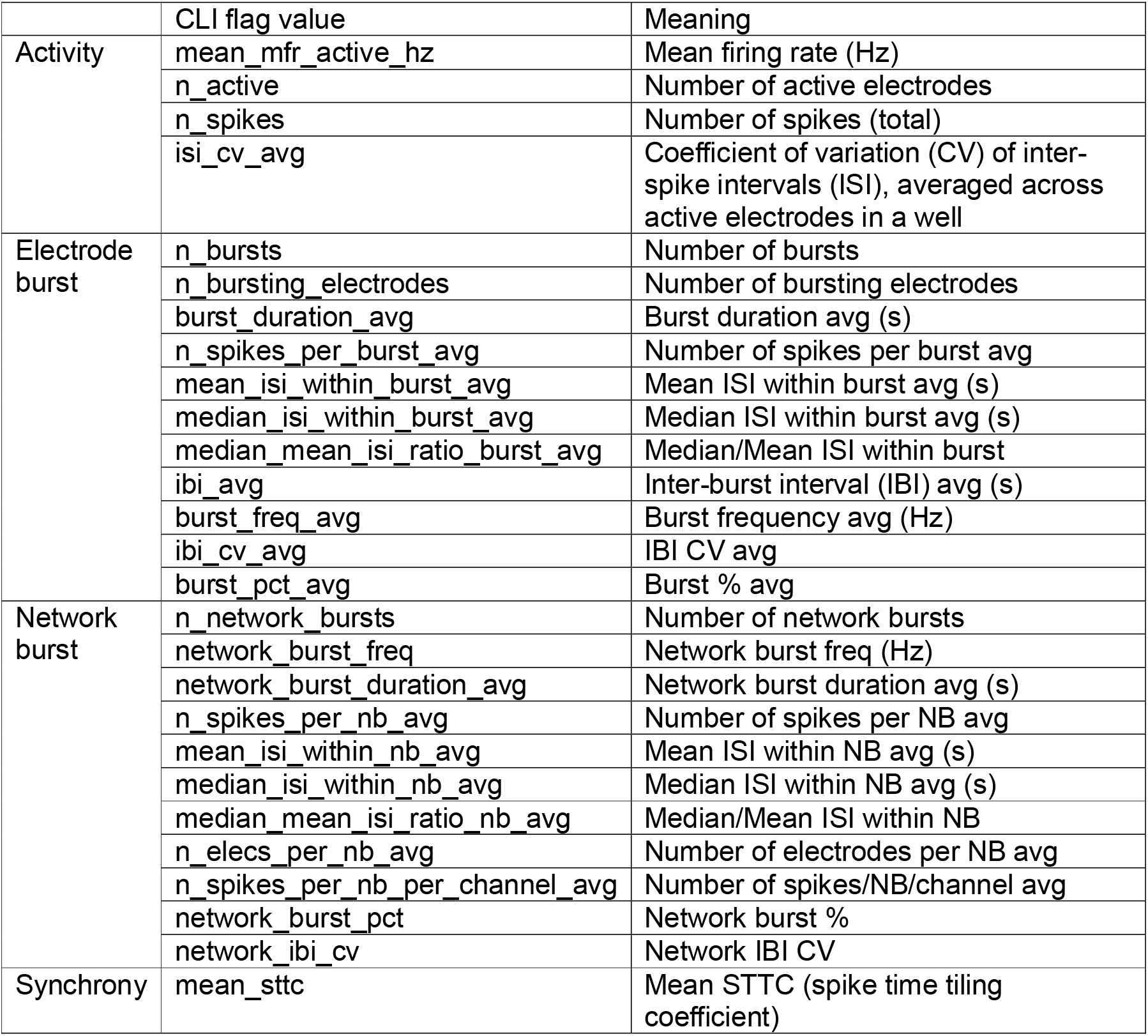
CLI flag values of metrics in py-mea-span.

As a proof of concept, we apply py-mea-span to a longitudinal CRISPR-mediated LGI2 knockdown experiment in iPSC-derived human cortical neurons. LGI proteins play important roles in synaptic transmission with LGI1 mutations underlying autosomal dominant epileptic disorders ^12^. LGI2, by contrast, remains understudied. Here, our MEA analysis reveals a reproducible network hyperexcitability phenotype upon LGI2 knockdown, suggesting that LGI2 also helps regulate neuronal network excitability.

## METHODS

### Cell line maintenance

Human CRISPRi-i3N stem cells carrying a stably integrated dCas9 construct were cultured at 37 °C and 5% CO_2_ in complete E8 medium (Gibco™), with daily medium changes. Cells were grown on Matrigel-coated plates (1:50 dilution in DMEM/F-12 HEPES; Gibco™). For passaging, cells were detached either with 0.5 mM EDTA and re-plated as colonies in E8 medium, or with Accutase and re-plated as single cells in E8 medium supplemented with 50 nM chroman-1 (Tocris).

Differentiation of stem cells into neurons was performed as previously described ^13^, with minor modifications. Following Accutase dissociation, 7 × 10^6^ or 1.5 × 10^7^ stem cells were seeded onto 10 or 15-cm Matrigel-coated dishes respectively. Cells were seeded in induction medium (IM), consisting of DMEM/F-12 HEPES supplemented with 1× N2, 1× NEAA, 1× GlutaMAX, and 2 μg/mL doxycycline. During initial plating, IM was additionally supplemented with 10 μM chroman-1, and the medium was replaced daily for the first three days.

Partially differentiated neurons were dissociated with Accutase and re-plated onto dishes coated with 100 μg/ml poly-L-ornithine (PLO) in cortical neuron (CN) medium, consisting of Neurobasal Plus supplemented with 1× B27, 10 ng/ml NT-3 (PeproTech), 10 ng/ml BDNF (PeproTech), and 1 μg/ml laminin. During re-plating, CN medium was supplemented with 1 μg/ml doxycycline. From this point onward, half-medium changes were performed three times per week.

### Generation of CRISPR knockdown cell lines

Guide RNAs targeting LGI2 were designed using the CRISPick server (Broad Institute; **Table 3**), selecting sites at or immediately upstream of the first exon. For targeting of a single gene, complementary oligonucleotides with appropriate overhangs were annealed and cloned into the BpiI site of the pKLV backbone (gift from the Prof. Evan Reid, Cambridge).

**Table 3.**
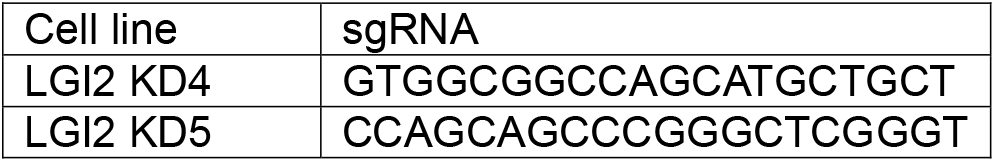
sgRNA used to generate LGI2 knockdown lines.

For lentiviral production, HEK293T cells were cultured in DMEM supplemented with 10% FBS at 37 °C and 5% CO_2_ and passaged twice weekly using Trypsin-EDTA. A total of 2 × 10^6^ cells were transfected in OptiMEM with 8 µl TransIT and 2 µg of plasmid DNA at a 3:2:1 ratio of CRISPRi pKLV construct:pCMVΔ8.91:pMD.G. The transfection mixture was added dropwise, and after 24 h the medium was replaced with E8. At 48 h post-transfection, the virus-containing E8 medium was collected, 0.45 μm filtered to remove debris and stored at - 70 °C.

5 × 10^5^ i3N stem cells were transduced with 2 ml of a 1:1 mixture of complete E8 medium and lentiviral supernatant supplemented with 50 nM chroman-1 and 10 µg/ml polybrene. Cells were seeded on Matrigel-coated 6-well plates and then transferred to 37 °C for incubation. After 24 h, the medium was replaced with fresh E8. At 48 h post-transduction, cells were switched to E8 medium containing 1 µg/ml puromycin, which was refreshed daily for at least 7 days. Surviving cells were subsequently maintained in puromycin-free E8 medium.

### Quantitative PCR (qPCR)

RNA was extracted from 2 × 10^6^ cells using the PureLink RNA extraction kit (Invitrogen) following the manufacturer’s protocol. cDNA was generated from purified RNA using the High-Capacity RNA-to-cDNA kit (Applied Biosystems) following the manufacturer’s protocol. qPCR reactions contained 1× TaqMan Fast Advanced Master Mix (Applied Biosystems), 50 ng cDNA, and 1× FAM-labelled TaqMan Gene Expression Assays (Applied Biosystems) specific to LGI2 (Hs00377867_m1). Reactions were run on a CFX96 Real-Time System (Bio-Rad), starting with 10 mins at 95 °C, followed by 40 cycles of 95 °C for 15 s and 60 °C for 1 min. Each experiment included n = 3 technical replicates per gene and N = 3 independent biological replicates. Gene expression changes were quantified using the ΔΔCt method with GAPDH as the reference gene. Statistical significance of fold changes was assessed by two-way ANOVA with Dunnett’s multiple comparisons testing.

### MEA recording

MEA recordings were conducted on cortical neurons cultured in 24-well CytoView MEA plates (Axion Biosystems, 16 electrodes per well) using the Maestro Edge multi-well plate recording platform (Axion Biosystems). Recordings were performed at 37 °C under 5% CO_2_. Action potentials were acquired continuously for 6 minutes per session, sampled at 12.5 kHz per electrode. The adaptive threshold for spike detection was set at 6× the standard deviation of the electrode noise. Recordings were conducted from DIV 12 to DIV 44 (DIV: days in vitro) across three independent biological replicates.

### Data processing

Raw spk files were generated by AxIS Navigator and were processed entirely using py-mea-span without prior conversion in NeuralMetric Tool. The following parameters were used: active electrode threshold, 5 spikes/min; burst detection, ISI-threshold algorithm (max_isi_s = 0.1 s, min_spikes = 5); network burst detection, combined-ISI algorithm (max_isi_s = 0.1 s, min_spikes = 50, participation_threshold = 0.35); STTC coincidence window, 50 ms. For each pair of electrodes A and B, T_A is the fraction of the recording that lies within ±Δt of any spike on electrode A. P_A is the fraction of spikes on electrode A that occur within ±Δt of a spike on electrode B. T_B and P_B are defined in the same way. The STTC is calculated as in Cutts & Eglen ^14^:

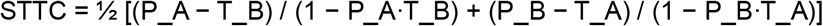

Pairwise comparisons between conditions at individual time points were performed on well-level metric values using Tukey’s Honest Significant Difference (HSD) test, with each well treated as an independent observation. NaN values from early DIVs from absent burst or network-burst events were treated as true zeros, reflecting absence of activity rather than missing data.

### Software implementation

py-mea-span is implemented in Python (version 3.10 or later) and depends only on widely available scientific libraries: numpy ^15^, scipy ^16^, pandas ^17^, matplotlib ^18^, pingouin ^19^, and statsmodels ^20^. The codebase is organised into six analysis modules. “io/” handles spk binary parsing and metadata loaders. “spike/” computes per-electrode firing-rate and inter-spike-interval metrics. “burst/” performs ISI-threshold burst detection and electrode-level burst statistics. “network/” performs combined-ISI network burst detection, network-burst metrics, and STTC synchrony. “stats/” exposes three pairwise condition-comparison tests through a unified pairwise_test dispatcher: Tukey’s Honest Significant Difference (parametric all-pairs, family-wise error controlled by the studentised-range distribution), pairwise Mann-Whitney U with Bonferroni correction, and Kruskal-Wallis with Dunn’s pairwise post-hoc with Bonferroni correction. As a guide, Tukey’s HSD is appropriate when values within each group are approximately normally distributed with similar variance. Kruskal-Wallis with Dunn’s post hoc test is the rank-based alternative for three or more groups when data are skewed or contain outliers; and Mann-Whitney U is the corresponding option for comparing two groups.

“viz/” produces spike rasters with array-wide spike-detection-rate histograms, ISI distributions, per-well electrode heatmaps, plate-layout heatmaps, pairwise STTC matrices, network-burst timelines, longitudinal trajectories, single-time-point violins with user-selectable significance brackets (any of the three pairwise tests, computed via the “stats/” dispatcher so that figure brackets and tabulated p-values are produced by identical code paths), and PCA scatter plots with optional loadings analysis. A pipeline.py module orchestrates the analysis flow, and a cli.py module provides the command-line entry point.

The package is distributed on PyPI (https://pypi.org/project/py-mea-span/), and version-controlled at https://github.com/Molecularbiologyworld/py-mea-span. The package includes 649 automated tests that check the spk file reader, the burst and network-burst detection, the metric-averaging behaviour, and every command-line option.

## RESULTS

### py-mea-span workflow and an LGI2 knockdown test case

py-mea-span implements the analysis workflow summarised in **Fig. 1A**. MEA recordings from Axion Biosystems CytoView 24-well plates (16 electrodes per well) are exported as binary spk files via AxIS Navigator. The package ingests the spk files and produces four classes of output: 1) spike raster plots show the timing of each spike on every electrode in a well, with an array-wide spike-detection rate (ASDR) histogram highlighting synchronised network bursts; 2) plate-layout heatmaps coloured by per-well metric values, giving an overview of the whole plate and revealing outliers or positional effects; 3) longitudinal time-course profiles across days in vitro, showing how activity matures and when conditions diverge; and 4) violin plots that compare the distribution of well-level values between conditions at a single time point.

**Figure 1.**
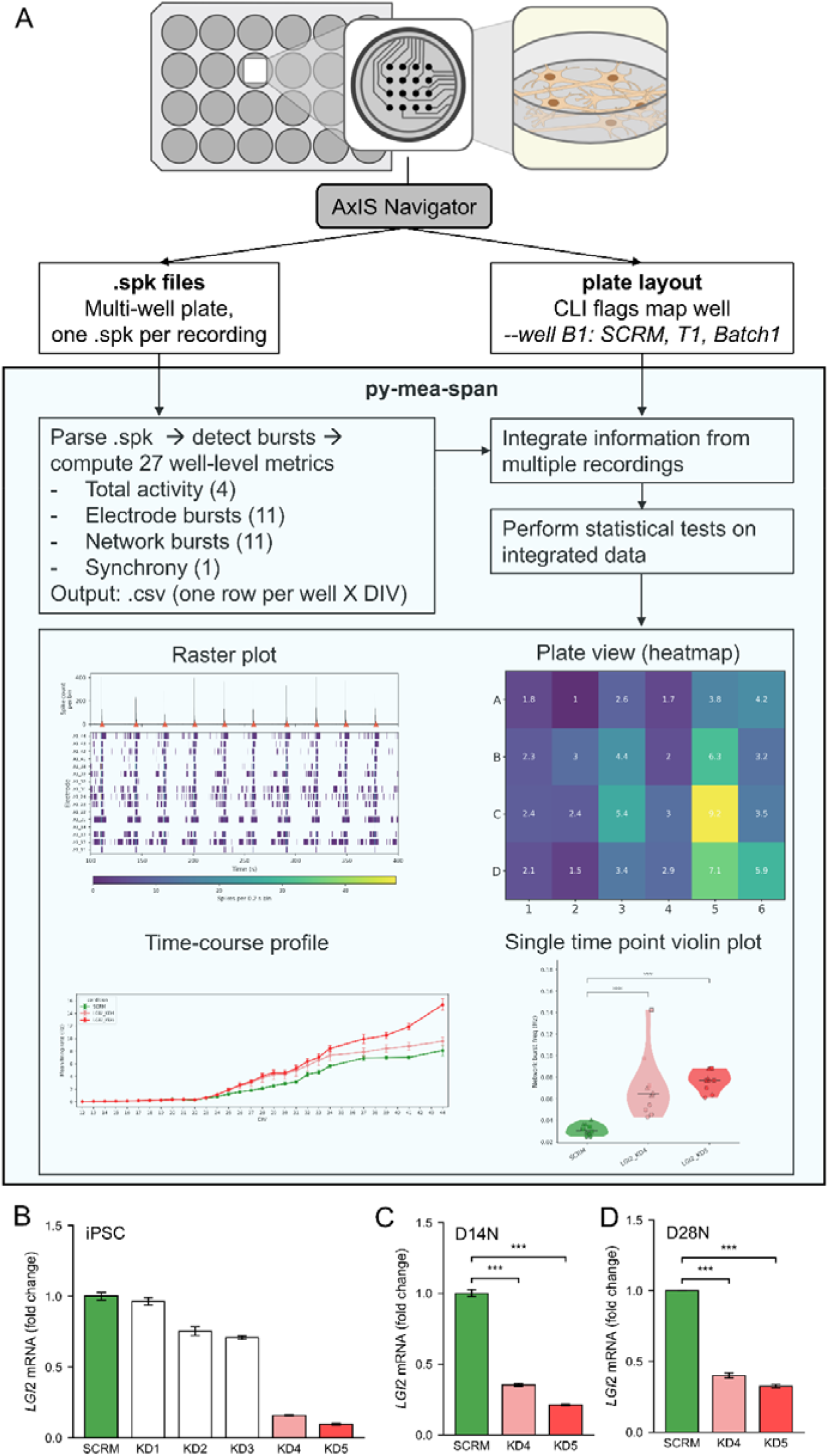
py-mea-span analysis pipeline for MEA recordings and LGI2 cell line validation. (A) Overview of the py-mea-span analysis workflow, with example outputs: raster plot of one well, plate heatmap, time-course profile of mean firing rate, and single time point violin plot of network burst frequency. (B-D) qPCR of LGI2 mRNA relative to scrambled control (SCRM) in stem cells at day 0 (B), neurons at day 14 (C) and neurons at day 28 (D). KD4 and KD5 were taken forward from the five knockdown lines. Bars show mean ± SEM, n = 3 technical replicates for (B-C), and N=3 biological replicates for (D). ***P < 0.001, one-way ANOVA with Dunnett’s multiple comparisons test.

To demonstrate the effectiveness and accuracy of the py-mea-span pipeline we tested it on a new CRISPR-mediated knockdown of Leucine-rich Glioma Inactivated 2 (LGI2) in iPSC-derived cortical neurons. In the LGI protein family, LGI2 is an under-investigated secreted protein predominantly expressed in neuronal tissues. In contrast to its paralog, LGI1, whose involvement in epilepsy has been confirmed in human patients ^21^ and transgenic mouse models ^22^, LGI2 remains a speculative candidate in human epilepsy. In canine models, secretion-deficient LGI2 mutations (causing LGI2 to be retained in the ER) have been shown to cause epilepsy ^23^. In mouse models, LGI2-deficient mice had no apparent neurological phenotype assessed by behavioural tests ^24^, yet a recent report showed that the hippocampal slices of LGI2 KO mice have the tendency to generate spontaneous ictal-like activity more often than wildtype slices ^25^. LGI2 was also among proteins reduced from the plasma membrane of neurons in a model of infantile epileptic disorder caused by loss of membrane gangliosides (GM3 synthase deficiency) ^26^.

We used CRISPRi-i3N induced iPSCs, which carry a doxycycline-inducible NGN2 transgene and a stably integrated catalytically inactive Cas9 for CRISPR interference ^13^. In this system, gene expression can be knocked down by introducing a single guide RNA (sgRNA) targeting the promoter of the gene of interest. Doxycycline-induced NGN2 expression drives rapid and uniform differentiation into cortical-like glutamatergic neurons. We define the day of doxycycline addition as day 0, and all time points are given as days in vitro (DIV) from induction. On day 3, neurons were detached and seeded onto MEA plates, and recordings were taken from DIV 12 to DIV 44.

Five independent CRISPRi knockdown lines were generated targeting LGI2. The two lines with the most substantial knockdown, LGI2 KD4 and KD5, were taken forward for analysis (**Fig. 1B**). qPCR confirmed a substantial reduction of LGI2 mRNA in both lines (**Fig. 1B&C**): 85-90% reduction at the stem cell stage, and 65-80% reduction by DIV 14. Because LGI2 loss-of-function is implicated in epilepsy, we hypothesised that LGI2 knockdown would alter spontaneous network activity, providing a biologically relevant test case for the package.

### py-mea-span reproduces NeuralMetric Tool outputs across all benchmarked metrics

To validate the quality of the outputs from py-mea-span we verified that it produces values matching those of the manufacturer’s NeuralMetric Tool (NMT) software, the reference point for Axion spk analysis. Across 946 well-observations spanning all three conditions and the full DIV 12-44 time window, the agreement was effectively perfect for every metric tested: mean firing rate (R^2^ = 1.0000; **Fig. 2A**), number of active electrodes (R^2^ = 1.0000; **Fig. 2B**), per-electrode burst frequency (R^2^ = 1.0000; **Fig. 2C**), and network burst frequency (R^2^ = 1.0000; **Fig. 2D**). The agreement spans the full dynamic range of each metric. The lower sample sizes for burst- and network-burst metrics reflect only well-observations in which the corresponding event class was detectable by either tool. py-mea-span therefore provides an open-source alternative to NMT, computing equivalent metrics directly from the same spk files.

**Figure 2.**
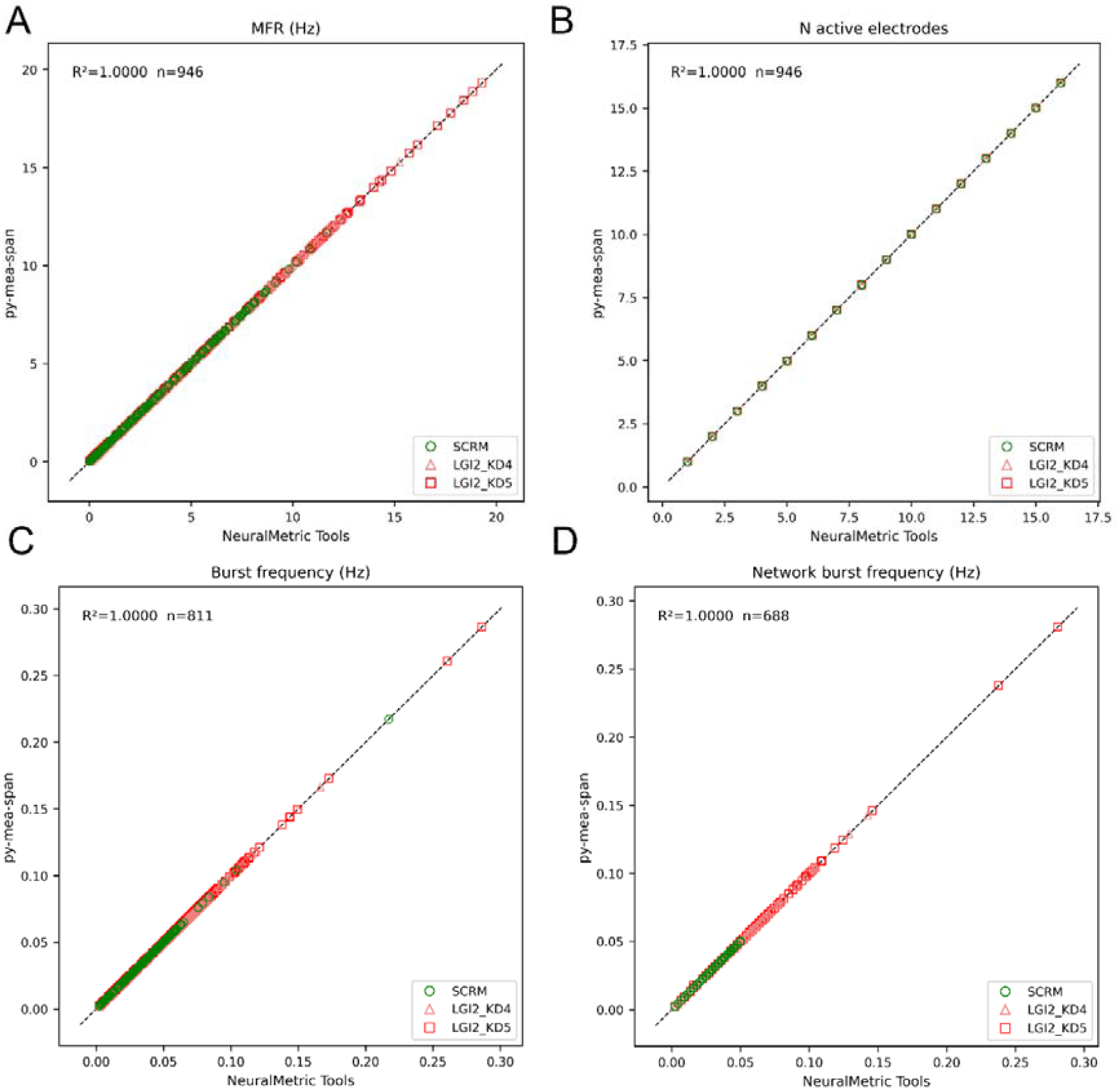
py-mea-span reproduces the metrics of the manufacturer’s NeuralMetric Tool. Four metrics computed by py-mea-span (y-axis) are plotted against the same metrics from NeuralMetric Tool (x-axis): (A) mean firing rate (MFR), (B) number (N) of active electrodes, (C) burst frequency and (D) network burst frequency. Each point is one well at one DIV, coloured by line (SCRM, LGI2 KD4, LGI2 KD5). The dashed line shows y = x. Coefficient of determination (R^2^) is calculated with all lines pooled.

### LGI2 knockdown produces a progressive network hyperexcitability phenotype from DIV 24 onward

We first compared spike raster plots from batch-matched SCRM and LGI2 KD5 neurons at DIV 28, after network activity had matured (**Fig. 3**). Both wells showed organised network bursting, with most electrodes firing together in bursts that appear as sharp peaks in the ASDR histogram (highlighted with red triangles). Interestingly, the LGI2 KD5 neurons produced more network bursts than the SCRM neurons over the same period.

**Figure 3.**
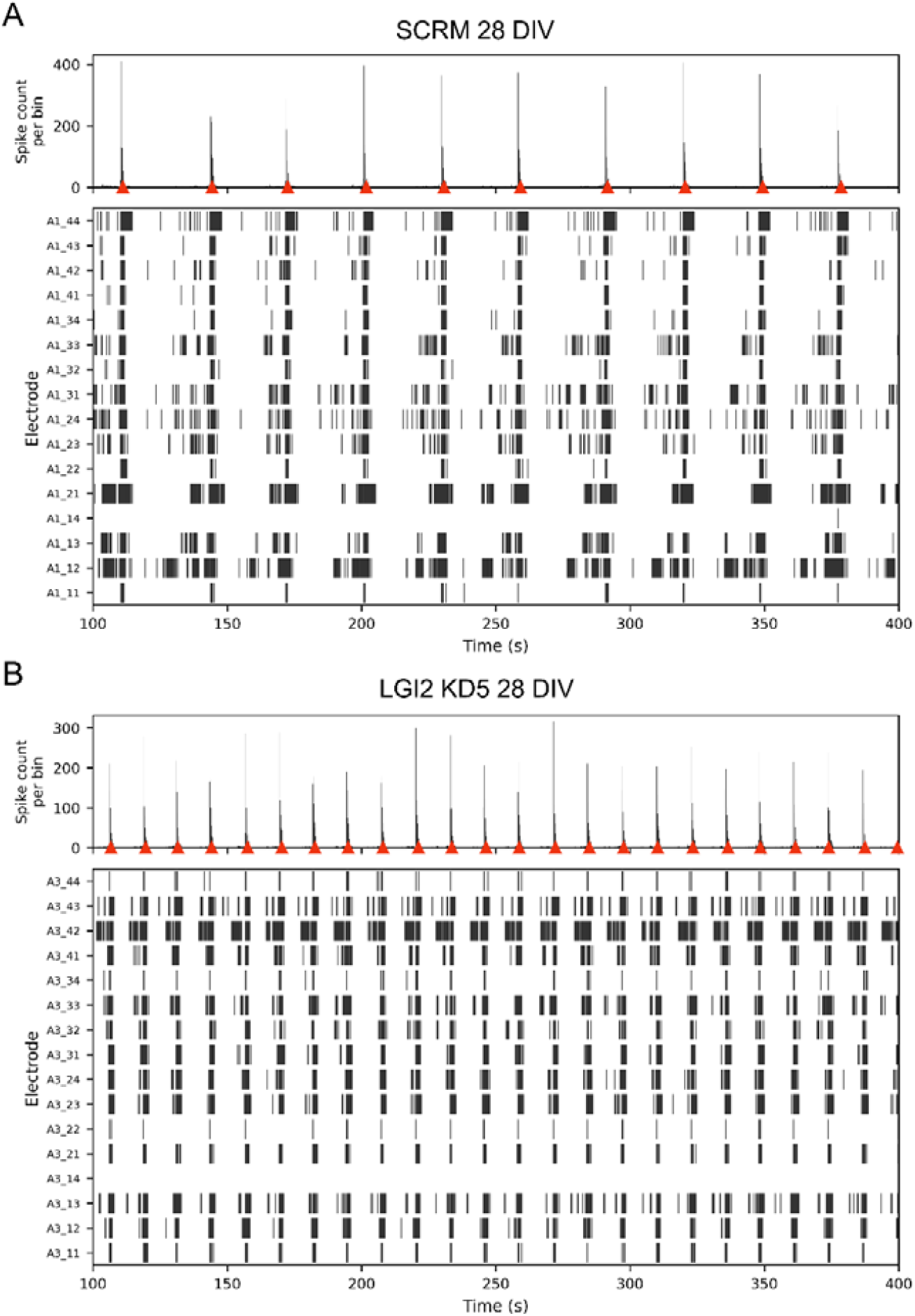
Spike rasters of representative SCRM (A) and LGI2 KD5 (B) wells at DIV 28. Top of each panel: array-wide spike-detection rate (ASDR, 0.2 s bins, 100-400 s); red triangles mark detected network bursts. Bottom: per-electrode raster plots in which each vertical tick is one detected spike and bursts appear as dense black blocks. Rows are the 16 electrodes of the well, labelled by well and electrode number as in Axion’s AxIS Navigator (e.g., A1_44 is electrode 44 of well A1).

Plate-layout heatmaps at DIV 28 (**Fig. 4**) showed that burst and network burst frequency were both higher in LGI2 KD4 and KD5 wells than in SCRM within every batch on both plates, so the effect was consistent across biological replicates.

**Figure 4.**
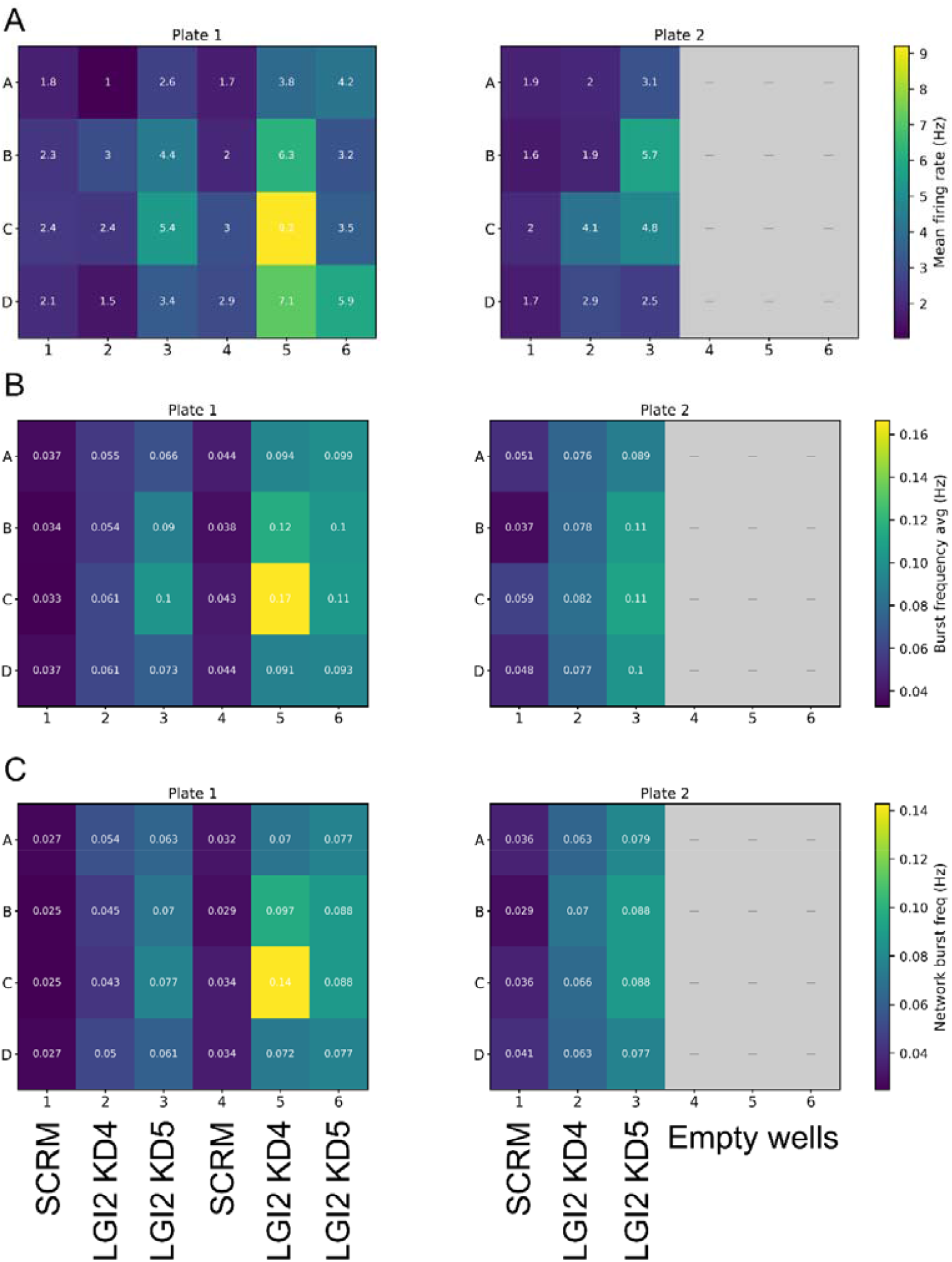
Plate-layout heatmaps of mean firing rate (A), per-electrode burst frequency (B), and network burst frequency (C) at DIV 28. Each panel shows two 24-well plates side-by-side; plate 2 columns 4-6 are unpopulated (grey). Columns 1/4 = SCRM, 2/5 = LGI2 KD4, 3/6 = LGI2 KD5. Each cell is annotated with its numerical value. Colour scale is set automatically from the lowest to the highest well value across both plates, so colours are comparable between plates.

The longitudinal time course showed when this phenotype emerges from DIV 24 (**Fig. 5**). Until DIV 23, all three lines developed similarly: the mean firing rates were low, network bursts first appeared around DIV 18, and at DIV 23 all cultures started to show consistent synchronised network bursting activities. Mean firing rate diverged gradually, most clearly in KD5 (**Fig. 5A**). This is consistent with the observation that KD5 line has a more effective knockdown of gene expression (**Fig. 1C**). From DIV 24 onward, both LGI2 KD lines diverged from SCRM. The divergence was abrupt for both burst and network burst frequency, which stayed elevated through DIV 44 (**Fig. 5B-C**).

**Figure 5.**
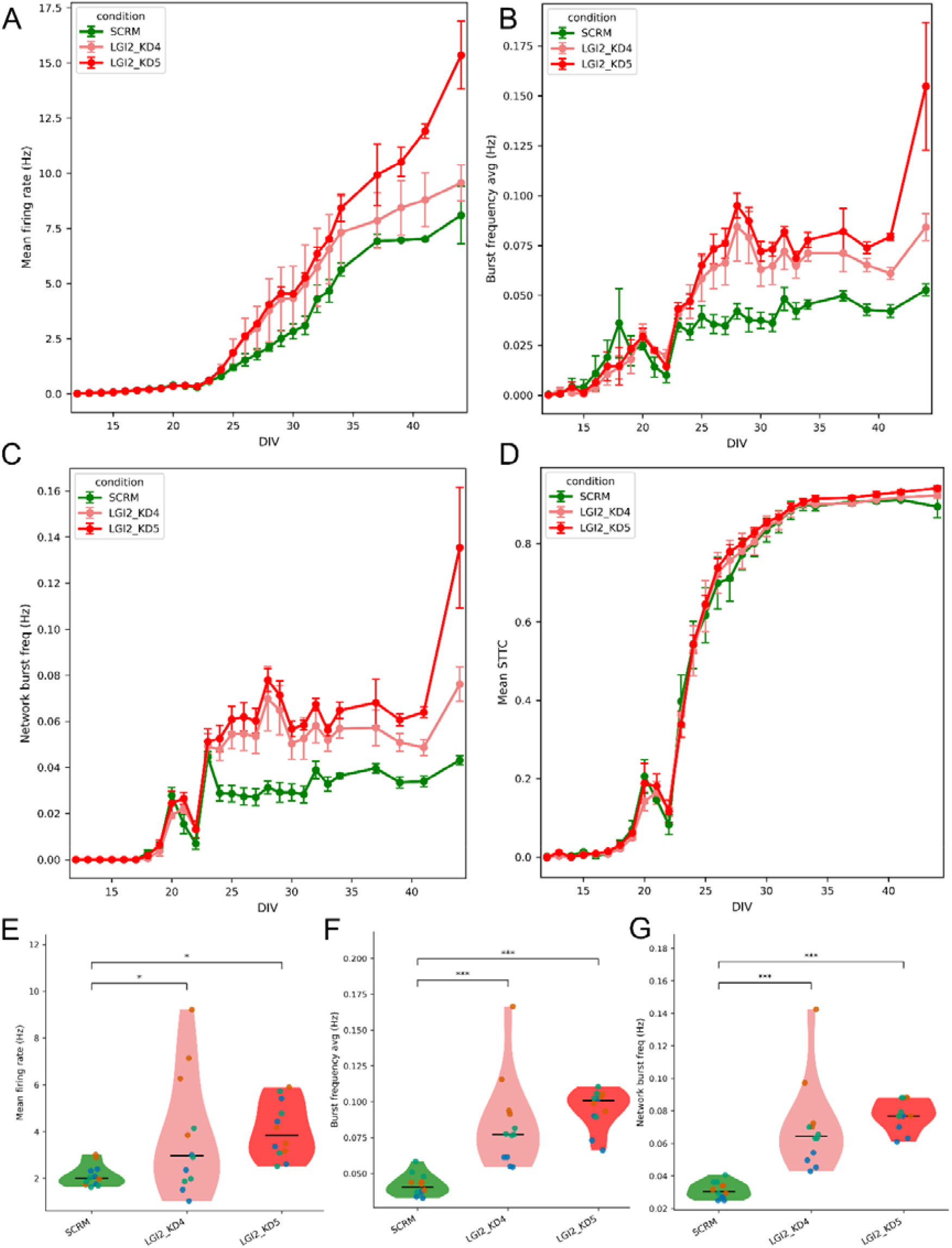
Longitudinal trajectories of network activity from DIV 12 to DIV 44 in SCRM LGI2 KD4 and LGI2 KD5 neurons. Each point is the mean ± SEM across n = 12 wells per condition. (A) Mean firing rate (Hz). (B) Mean per-electrode burst frequency (Hz). (C) Network burst frequency (Hz). (D) Mean spike-time tiling coefficient (STTC). (E-G) Mean firing rate, burst frequency and network burst frequency at DIV 28 across the three conditions (n = 12 wells/condition). * p < 0.05, *** p < 0.001, Tukey HSD-adjusted p-values.

In contrast, synchrony did not differ between these lines. We measured synchrony with the spike time tiling coefficient (STTC). STTC quantifies how often spikes on one electrode fall within a short time window of spikes on another, relative to the coincidences expected by chance. It ranges from -1 to 1, where 1 indicates perfectly synchronised spiking and 0 indicates chance-level coincidence, and we averaged it over all pairs of active electrodes in each well. We chose STTC because, unlike the commonly used correlation index, it is largely independent of firing rate ^14^. This matters here because the knockdown lines fire and burst more than SCRM, so a rate-dependent measure could report higher synchrony simply because there are more spikes. Mean STTC rose in parallel in all three lines and plateaued around 0.9 from DIV 33 (**Fig. 5D**). It did not differ between lines even from DIV 24 to 30, before it approached its ceiling. This indicates that LGI2 knockdown elevates network excitability without altering the temporal structure of multi-electrode synchrony.

To compare the lines statistically at a single time point, we plotted the DIV 28 values as violin plots (Fig. 5E-G), which show the full distribution of all wells. At DIV 28, both knockdown lines had significantly higher burst and network burst frequencies than SCRM (**Fig. 5F-G**). The increase in mean firing rate was smaller and more variable (**Fig. 5E**). py-mea-span offers three statistical tests: Tukey’s HSD, Mann-Whitney U, and Kruskal-Wallis with Dunn’s post hoc test.

## DISCUSSION

py-mea-span provides an open-source, end-to-end Python pipeline for Axion Biosystems multi-electrode array recordings. It reads the proprietary spk binary format directly. From these files it calculates per-electrode and network-level metrics, runs statistical comparisons, and produces publication-ready figures. It can be used from either a Python API or a mea-span command-line interface. Across 946 well-observations the package reproduced the manufacturer’s NeuralMetric Tool reference at R^2^ = 1.0000 for every benchmarked metric.

In addition to reproducing the accuracy of NMT, it offers more flexibility as most parts of the analysis can be changed (**Table 4**). Users can define the analysis to part of a recording, or to selected wells, conditions and days in vitro. They can adjust the thresholds for detecting bursts and network bursts, and the time window used to measure synchrony. Users can also choose suitable statistical test, and decide whether wells or biological replicates count as independent samples. Plot appearance, such as colours, marker shapes and figure size, are set with simple command-line flags.

**Table 4.** Customizable features of py-mea-span.

| Feature | What can be changed | Command-line flags |
| --- | --- | --- |
| Recording time window | Analyse only part of a recording | --time-start, --time-end |
| Time points | Individual DIVs or a DIV range | --div, --div-min, --div-max |
| Wells, conditions, batches | Which wells are analysed; which conditions are plotted and in what order; filtering by any column | --wells, --group-order, --filter |
| Metrics | Any of the 27 well-level metrics | --metric |
| Active electrodes | Minimum firing rate for an electrode to count as active | --active-threshold |
| Bursts | Maximum inter-spike interval and minimum spikes per burst | --max-isi, --min-spikes |
| Synchrony | STTC coincidence window | --sttc-dt |
| Statistics | Test (Tukey's HSD, Mann–Whitney U, or Kruskal–Wallis with Dunn's), pairs compared, wells or biological replicates as the unit of analysis | --test, --compare, --pool |
| Plot appearance | Condition colours; point colour, shape and size by any column (e.g. batch); figure size; heatmap colour map; raster histogram bin width | --color, --point-hue, --point-color, --point-shape, --point-shape-map, --point-size, --figsize, --cmap, --asdr-bin |
| Plate layout | Condition, biological and technical replicate, and batch for each well; reusing a saved layout | --well, --layout-from |

Beyond demonstrating the package, the LGI2 KD neuronal recordings contribute biologically meaningful information. These data using human iPSC-derived-neurons demonstrate that LGI2 deficiency drives network-level changes in spontaneous electrical activity. The reproducible elevation of firing, burst, and network-burst rates emerging from DIV 24 onward, in both knockdown lines, may be consistent with the epilepsy-related phenotypes previously reported in canine and mouse LGI2 loss-of-function models ^23,25^. Our observation thus extends the LGI2-related epilepsy literature into a tractable human cellular system.

By bringing both raw spk file processing and statistical analyses into one open-source Python environment, py-mea-span fills a gap not addressed by existing open-source MEA tools, which require pre-processing through proprietary software, and complements the manufacturer’s pipeline by adding statistical tests. This package extends previously available analytical resources for MEA data analysis and we anticipate future extensions of this with the expanding use of this technique.

## ACKNOWLEDGEMENTS

Large language models were used to assist with code optimization. The funders had no role in study design, data collection and analysis, decision to publish, or preparation of the manuscript. For the purpose of open access, the author has applied a Creative Commons Attribution (CC BY) license to any Author Accepted Manuscript version arising from this submission.

## REFERENCES

1. Cotterill, E. et al. Characterization of Early Cortical Neural Network Development in Multiwell Microelectrode Array Plates. J. Biomol. Screen. 21, 510–519 (2016).

2. Olivero-Acosta, M. I. et al. Epilepsy-Associated SCN2A-L1342P Mutation Drives Network Hyperexcitability and Widespread Transcriptomic Changes in Human Cortical Organoids. Preprint at 10.1101/2025.08.18.670956 (2025).

3. Wainger, B. J. et al. Intrinsic membrane hyperexcitability of amyotrophic lateral sclerosis patient-derived motor neurons. Cell Rep. 7, 1–11 (2014).

4. Weigel, B. et al. MYT1L haploinsufficiency in human neurons and mice causes autism-associated phenotypes that can be reversed by genetic and pharmacologic intervention. Mol. Psychiatry 28, 2122–2135 (2023).

5. Fan, J. et al. Assessing seizure liability using multi-electrode arrays (MEA). Toxicol. Vitro Int. J. Publ. Assoc. BIBRA 55, 93–100 (2019).

6. McConnell, E. R., McClain, M. A., Ross, J., Lefew, W. R. & Shafer, T. J. Evaluation of multi-well microelectrode arrays for neurotoxicity screening using a chemical training set. Neurotoxicology 33, 1048–1057 (2012).

7. Habela, C. W. et al. Altered development and network connectivity in a human neuronal model of 15q11.2 deletion-related neurodevelopmental disorders. Transl. Psychiatry 15, 329 (2025).

8. Trujillo, C. A. et al. Complex Oscillatory Waves Emerging from Cortical Organoids Model Early Human Brain Network Development. Cell Stem Cell 25, 558–569.e7 (2019).

9. Gordon, E. A., Bennett, D. L., Baskozos, G. & Comini, M. MEAanalysis: an open-source R package for downstream visualization of AxIS navigator multi-electrode array burst data at the single-electrode level. Bioinforma. Adv. 5, vbaf160 (2025).

10. Gelfman, S. et al. meaRtools: An R package for the analysis of neuronal networks recorded on microelectrode arrays. PLoS Comput. Biol. 14, e1006506 (2018).

11. Hu, M. et al. MEA-ToolBox: an Open Source Toolbox for Standardized Analysis of Multi-Electrode Array Data. Neuroinformatics 20, 1077–1092 (2022).

12. Kegel, L., Aunin, E., Meijer, D. & Bermingham, J. R. LGI proteins in the nervous system. ASN Neuro 5, 167–181 (2013).

13. Fernandopulle, M. S. et al. Transcription Factor–Mediated Differentiation of Human iPSCs into Neurons. Curr. Protoc. Cell Biol. 79, (2018).

14. Cutts, C. S. & Eglen, S. J. Detecting pairwise correlations in spike trains: an objective comparison of methods and application to the study of retinal waves. J. Neurosci. Off. J. Soc. Neurosci. 34, 14288–14303 (2014).

15. Harris, C. R. et al. Array programming with NumPy. Nature 585, 357–362 (2020).

16. Virtanen, P. et al. SciPy 1.0: fundamental algorithms for scientific computing in Python. Nat. Methods 17, 261–272 (2020).

17. McKinney, W. Data Structures for Statistical Computing in Python. in 56–61 (Austin, Texas, 2010). doi:10.25080/Majora-92bf1922-00a.

18. Hunter, J. D. Matplotlib: A 2D Graphics Environment. Comput. Sci. Eng. 9, 90–95 (2007).

19. Vallat, R. Pingouin: statistics in Python. J. Open Source Softw. 3, 1026 (2018).

20. Seabold, S. & Perktold, J. Statsmodels: Econometric and Statistical Modeling with Python. in 92–96 (Austin, Texas, 2010). doi:10.25080/Majora-92bf1922-011.

21. Ottman, R. et al. LGI1 mutations in autosomal dominant partial epilepsy with auditory features. Neurology 62, 1120–1126 (2004).

22. Fukata, Y. et al. Disruption of LGI1-linked synaptic complex causes abnormal synaptic transmission and epilepsy. Proc. Natl. Acad. Sci. U. S. A. 107, 3799–3804 (2010).

23. Seppälä, E. H. et al. LGI2 truncation causes a remitting focal epilepsy in dogs. PLoS Genet. 7, e1002194 (2011).

24. Kozar-Gillan, N. et al. LGI3/2-ADAM23 interactions cluster Kv1 channels in myelinated axons to regulate refractory period. J. Cell Biol. 222, e202211031 (2023).

25. Savonlehto, T. et al. Lgi2-deficient mice manifest epileptiform activity in the developing hippocampal network and ADHD-like behavioural comorbidity in adulthood. Exp. Neurol. 396, 115541 (2026).

26. Barrow, H. G. et al. Sialylated Gangliosides are Required for Plasma Membrane Organization and Neuronal Function. Preprint at 10.64898/2026.03.18.712603 (2026).

